# A proton pump enhancing photosynthesis links phagocytosis to marine algae symbiogenesis

**DOI:** 10.1101/2022.05.26.493626

**Authors:** Daniel P. Yee, Ty J. Samo, Raffaela M. Abbriano, Bethany Shimasaki, Maria Vernet, Xavier Mayali, Peter K. Weber, B. Greg Mitchell, Mark Hildebrand, Martin Tresguerres

## Abstract

Diatoms, dinoflagellates, and coccolithophorids are the dominant groups of marine eukaryotic phytoplankton collectively responsible for the majority of primary production in the ocean^1^. These phytoplankton contain additional intracellular membranes around their chloroplasts derived from ancestral engulfment of red microalgae by unicellular heterotrophic eukaryotes that led to secondary endosymbiosis^2^. This symbiogenesis hypothesis for the origin of modern secondary endosymbiotic phytoplankton is supported by a wealth of palaeontologic, morphologic, and genomic evidence^3–6^. However, the selectable evolutionary advantage of these membranes and the physiological significance for extant phytoplankton are unknown. We report that the proton-pumping enzyme V-type H^+^-ATPase (VHA), ubiquitously used in eukaryotic intercellular digestion, is localized around the chloroplasts of centric diatoms and that VHA-activity significantly enhances photosynthesis over a wide range of oceanic irradiances. Similar results in pennate diatoms, dinoflagellates, and coccolithophorids, but not green or red microalgae, imply a mechanism resulting from the co-option of phagocytic VHA activity into a carbon concentrating mechanism that is common to secondary endosymbiotic phytoplankton. Furthermore, analogous VHA-dependent mechanisms in extant photosymbiotic marine invertebrates^7–9^ provide functional evidence for an adaptive advantage throughout the transition from endosymbiosis to symbiogenesis. Our results suggest that VHA-dependent enhancement of photosynthesis contributes at least 7% of primary production in the ocean, providing an example of a symbiosis-derived evolutionary innovation with global environmental implications.

## Main Text

Symbiogenesis refers to the fusing of two organisms into a new single organism with emerging evolutionary innovations^3,10^. Diatoms, dinoflagellates, and coccolithophorids are unicellular phytoplankton that originated from the phagocytosis of a red alga by a heterotrophic protozoan and, over evolutionary time, culminated in their fusion into single organisms carrying ancestral red chloroplasts^11^. Today, these secondary endosymbiotic phytoplankton dominate our oceans and are collectively responsible for the majority of the primary production in the ocean^1^.

Secondary endosymbiotic phytoplankton possess additional intracellular membranes surrounding their chloroplasts, which are hypothesized to derive from the ancestral phagosome that engulfed the red alga^2^. Since intracellular digestive vacuoles are ubiquitously acidified by V-type H^+^-ATPase (VHA) proton pumps^12^, acidification of the microenvironment around secondary chloroplasts was proposed to promote the dehydration of dissolved inorganic carbon (DIC) into CO_2_ thus enhancing photosynthesis^13,14^. Evidence for VHA-enhancement of photosynthesis in phytoplankton has yet to be reported; however, an analogous mechanism has been recently identified in cnidarian and mollusks that establish tertiary phago-photosymbiotic relationships with microalgae^7–9^. Here, we investigated whether VHA activity enhances photosynthetic O_2_ production by extant marine diatoms, dinoflagellates and coccolithophorids, and conducted detailed experiments on the diatom *Thalassiosira pseudonana* to confirm the presence of VHA surrounding their chloroplasts and to quantify the contribution of VHA activity to photosynthetic carbon fixation over the full range of environmentally relevant irradiances.

Similar to photosymbiotic cnidarians and mollusks^7–9^, inhibition of VHA activity induced significant decreases in gross maximum O_2_ production in the centric diatom *T. pseudonana*, the pennate diatom *Phaeodactylum tricornutum*, the dinoflagellate *Brandtodinium nutricula*, and the coccolithophorid *Emiliania huxleyi*. In contrast, photosynthetic O_2_ production by the green alga *Chlorella protothecoides* or by the red alga *Porphyrydium purpurneum* were not affected by VHA inhibition (Fig. 1; Extended Data Table 1). While these latter two species possess VHA that contribute to vacuole homeostasis^15–18^, they lack the intracellular membranes of phagocytic origin that surround the chloroplasts of secondary endosymbiotic phytoplankton.

**Figure 1.**
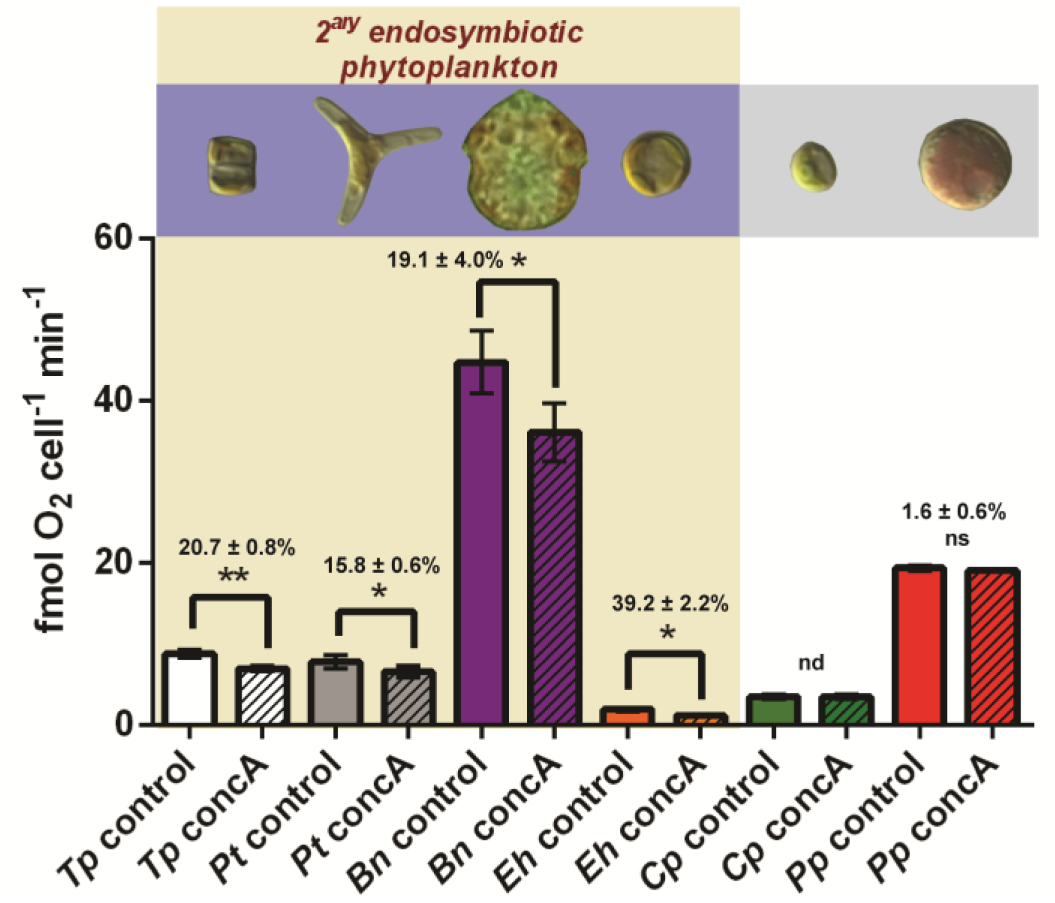
Contribution of VHA to O_2_ production of marine microalgae. Gross maximum O_2_ production per cell in control (vehicle DMSO; open bars) and VHA-inhibited (concA=10 nM concanamycin A; hatched bars) cultures. Percent reduction of O_2_ production and SEM between control and VHA-inhibited cultures are displayed above paired data bars. *Tp= T. pseudonana*(centric diatom); *Pt= P. tricornutum* (pennate diatom); *Bn= Brandtodinium nutricula* (dinoflagellate); *Eh= E. huxleyi* (coccolitophorid); *Cp= C. protothecoides* (green microalgae); *Pp=P. purpurneum* (red microalgae). Error bars = SEM; n = 3 except *Br* (n=6); Paired *t* test: \**p<0*.05; \*\**p<0*.01; nd= no difference; ns=no significant difference.

We further explored VHA-enhancement of photosynthesis in the model diatom *T. pseudonana*.Transcriptomics analysis on cell-cycle synchronized cultures revealed constitutive expression of all VHA subunits (Extended Data Fig. 1a). However, VHA can have multiple subcellular localizations and participate in diverse processes in addition to intracellular digestion^19,20^. Indeed, we recently showed that VHA localizes to the silica deposition vesicle of *T. pseudonana* during the G2+M phase of cellular division, and that VHA activity is essential for biomineralization of the silica cell wall^21^. Here, 3D confocal microscopy of transgenic *T. pseudonana* expressing eGFP-tagged VHA subunit B demonstrated that this proton pump is present around chloroplasts throughout the cell cycle (Fig. 2 & Extended Data Fig. 1c). Simultaneous accumulation of the acidotropic fluorescent dye PDMPO^22^ around chloroplasts implied acidification of the microenvironment to ≤ pH 5.5. At this pH, the majority of DIC exists as CO_2_, supporting the quarter-century-old hypothesis that intracellular secondary endosymbiotic membranes play a role in enhancing photosynthesis^13,14^.

**Figure 2.**
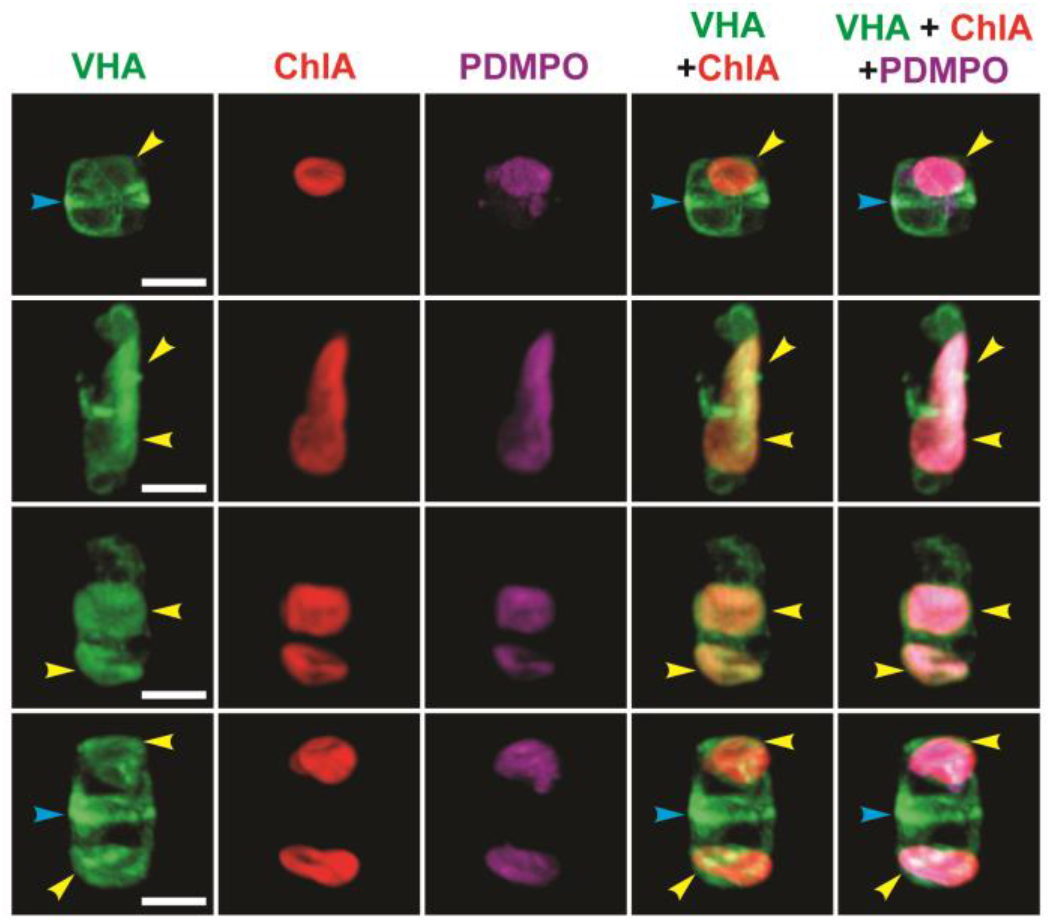
VHA surrounds the chloroplasts of *T. pseudonana*. 3D confocal images of eGFP-tagged VHA_B_ around chloroplasts (yellow arrows) co-localized with the acidotrophic dye PDMPO (magenta), and silica deposition vesicles (blue arrows) at different cell-cycle stages [scale bars: 5 μm].

We used NanoSIMS to quantify the effect of VHA inhibition on photosynthetic carbon fixation and incorporation into individual *T. pseudonana* cells following an 8h incubation with NaH^13^CO_3_. The isotope images revealed a statistically significant ~50% decrease in net biomass formation from photosynthesis in VHA-inhibited diatoms compared to controls (Fig. 3a).

**Figure 3.**
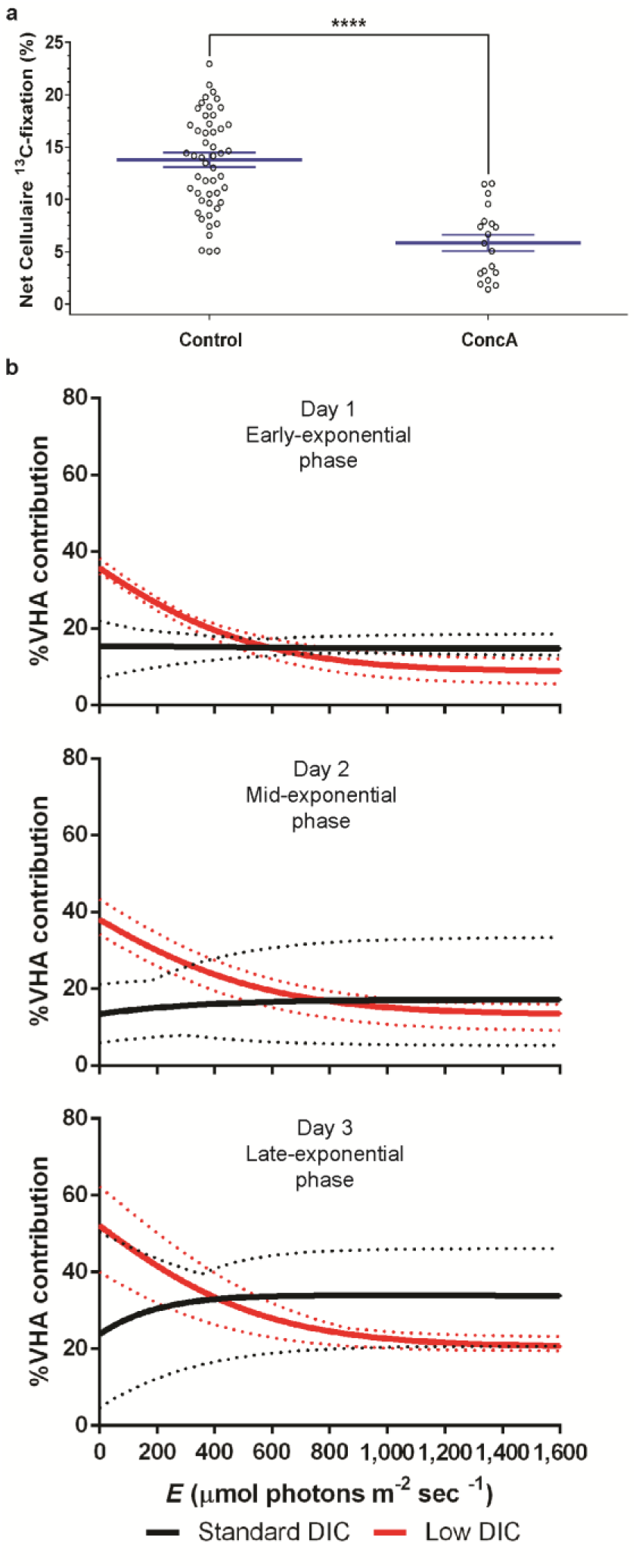
Contribution of VHA to carbon fixation in *T. pseudonana*. (**a**) Single-cell NanoSIMS quantification of net biomass formation from photosynthesis in controls (n=51) and VHA-inhibited (concA= 10 nM concanamycin A) cells (n=19). Error bars = SEM; Paired *t* test: \*\*\*\**p*<0.0001. (**b**) Contribution of VHA to bulk ^14^C incorporation over the full range of oceanic irradiances, at three stages of growth, and at standard (1.92 mM; black lines; n=6) and low (1.60 mM; red lines; n=3) seawater DIC levels (dotted lines= 95% CI).

We conducted a more complete assessment of VHA-dependent photosynthetic carbon fixation by quantifying ^14^C incorporation into *T. pseudonana* following 1h incubations with NaH^14^CO_3_. To capture the varying physiological status of cells during a diatom bloom^23^, measurements were taken during early-, mid-, and late-exponential growth phases over three days of culturing (Extended Data Fig. 3). To assess the broad range of oceanic irradiances resulting from latitude, clouding, depth, and ocean mixing^24^, photosynthesis versus irradiance (P-E) incubations were curve fitted from 0-1,600 μmols photons m^-2^ sec^-1^. And to examine the response to rapid environmental DIC changes, incubations were conducted at ~2.0 mM (standard) and ~1.6 mM (low) DIC encompassing levels in the open ocean and coastal environments due to rainfall, ice melting, and terrestrial freshwater inputs^25–27^ (Extended Data Fig. 4).

In our experiments, VHA inhibition impaired carbon fixation by at least 13.5% and as much as 52%. At standard DIC, the contribution of VHA to carbon fixation was relatively constant across irradiances ranging from ~15% on day 1 to ~29% on day 3 (Fig. 3b; black curves). At low DIC, VHA contribution also increased from day 1 to 3; however, it gained additional importance at irradiances under 500 μmols photons m^-2^sec^-1^. At sub-saturating irradiances <100 μmols photons m^-2^ sec^-1^, the contribution of VHA to carbon fixation approached 40% on days 1-2 and surpassed 50% on day 3 of culturing (Fig. 3b; red curves). This pattern implies that VHA-enhancement of photosynthesis is most significant during DIC limitation and under light levels that match ocean depths where phytoplankton are most abundant^28^.

In combination, the O_2_ production, ^13^C-NanoSIMS, and ^14^C-P-E measurements demonstrate that VHA activity significantly contributes to photosynthetic fixing of carbon that is retained as biomass. Given that diatoms contribute nearly 50% of carbon fixation in the ocean^29,30^, VHA-enhanced photosynthesis can be estimated to contribute between ~7 and 25% of oceanic primary production, or between 3.5 and 13.5 Gtons of fixed carbon per year (Extended Data Table 3). These numbers can only increase after accounting for analogous VHA-dependent mechanisms in other secondary endosymbiotic phytoplankton (Fig. 1) and tertiary photosymbiotic invertebrates^7–9^.

Engulfment of food particles by phagocytosis followed by lysosomal digestion is ubiquitously used by eukaryotic cells^12^, protozoans^11^, and invertebrate animals^31^. Acidification by VHA-which is conserved in all eukaryotes-is essential to these processes^20^. The presence of VHA in membranes of phagocytic origin surrounding diatom chloroplasts and cnidarian endosymbiotic microalgae and its role in enhancing photosynthesis provide a functional link between phagocytosis, endosymbiosis and symbiogenesis. The symbiotic microalgae of giant clams are hosted in the gut lumen and therefore are extracellular; however, they are also surrounded by a host-derived VHA-containing membrane whose primary purpose is food digestion. Thus, VHA activity at the symbiosis interface constitutes a broader mechanism to enhance photosynthesis in phago-photosymbioses (Fig. 4). Acidification of the microenvironment surrounding the chloroplast or microalgae is bound to promote CO_2_ accumulation, prevent CO_2_ back-flow into the cytoplasm, and ultimately help saturate RuBisCO with CO_2_ thus maximizing carbon fixation rates^14,32^. Interestingly, the carbon concentrating mechanisms of diatoms, corals, and giant clams^33–35^ all rely on carbonic anhydrases, and these enzymes form ubiquitously functional complexes with VHA to acidify intra- and extra-cellular compartments for diverse functions^19^ including lysosomal^36^ and epithelial digestion^37^.

**Figure 4.**
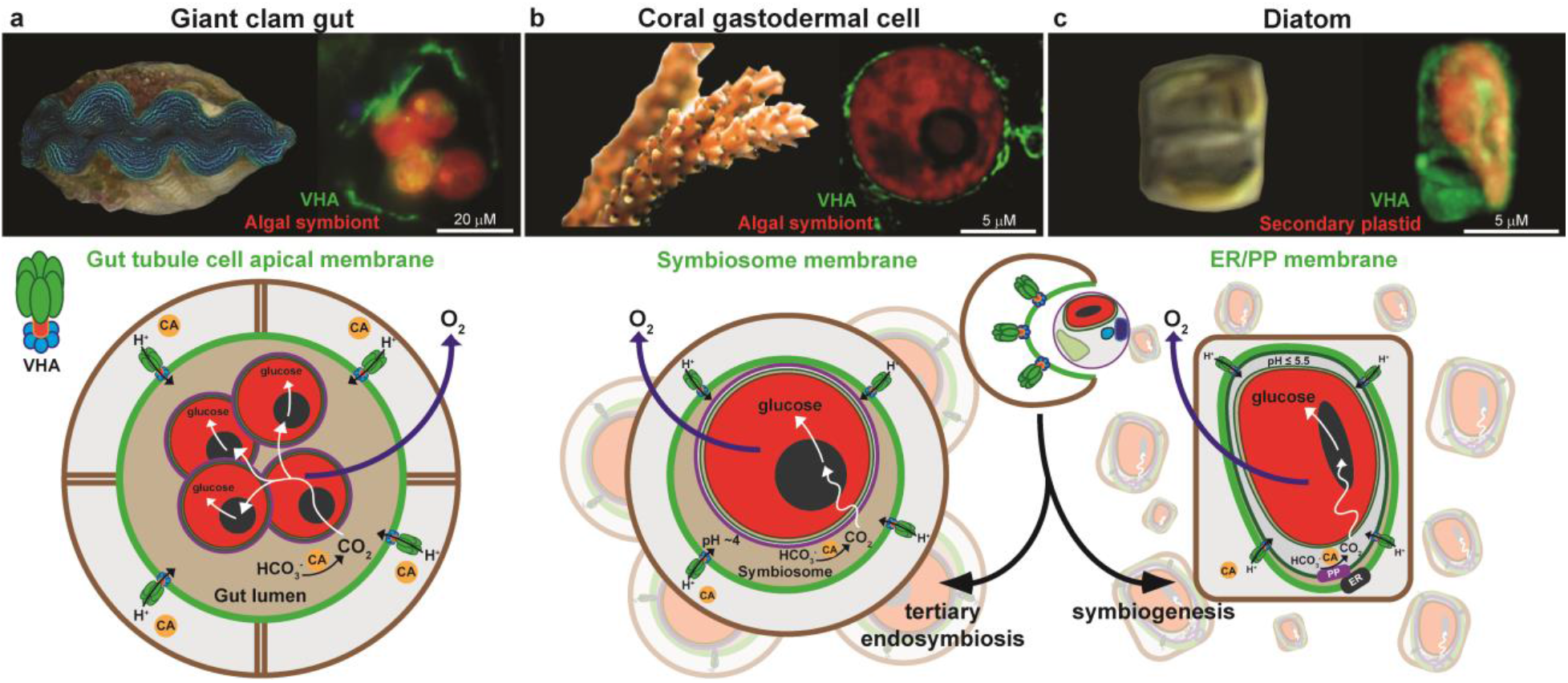
Enhancement of photosynthesis by VHA in diverse marine phago-photosymbioses. VHA in the biological membrane at the photosymbiosis interface promotes photosynthesis at diverse stages of phago-photosymbiotic integration. (**a**) the apical membrane of epithelial cells in giant clam gut tubules that host microalgae extracellularly^8^, (**b**) the symbiosomal membrane of coral gastrodermal cells that host microalgae intracellularly^7^, and (**c**) the endoplasmic reticulum/periplastid (ER/PP) membranes of diatoms that surround the plastid acquired from red microalgae by phagocytosis.

VHA-enhancement of photosynthesis co-opted from intracellular digestion confers a clear ecological advantage to extant marine secondary endosymbiotic phytoplankton. Since this advantage is also evident in extant tertiary photosymbiotic invertebrates, we speculate it was similarly important for the ancestral heterotrophic protist(s) that engulfed red microalgae during the transitional stages towards symbiogenesis. VHA-enhancement of photosynthesis would have provided a trait for positive selection and evolutionary advantage over other phytoplankton, particularly in the low-CO_2_ Permian oceans where diatoms originated^13,14^.

## Materials and Methods

### Culturing Conditions

Cultures of *T. pseudonana* (CCMP1335), *T. pseudonana* expressing eGFP-tagged VHA_B_^21^, *P. tricornutum* (CCMP632), *E. huxleyi* (CCMP1516), *P. purpurneum* (CCMP1947), and *C. protothecoides* (CCAP-211/7a) were grown axenically in F/2 medium prepared with seawater from the Scripps Pier. Inoculum cultures were maintained in 50 ml volumes in 125 ml Erlenmeyer flasks on an orbital shaker under continuous illumination provided by cool-white fluorescent lamps at 70 μmol photons m^-2^ s^-1^ at 18°C. Inoculum grown to ~2×10^6^ cells ml^-1^ were used to inoculate 1 l bioreactor tubes bubbled with air and under continuous illumination by cool-white fluorescent lamps at ~100 μmol photons m^-2^ s^-1^ at 18°C at a starting density of 1×10^5^ cells ml^-1^. Cultures of *B. nutricula* (RCC3387) were grown in T25 tissue culture flasks (Falcon) with 25 ml of K2 medium prepared with seawater collected from Villefranche-sur-mer Marine Station. Cultures were inoculated at 4×10^4^ cells ml^-1^ maintained at 70 μmol photons m^-2^ s^-1^ at 20°C without bubbling or agitation.

Daily measurements of cell counts, chlorophyll content, pH, and total CO_2_ (TCO_2_) were collected over the duration of all culturing experiments except for *B. nutricula* in which only cell counts and chlorophyll content was collected. These measurements were used to calculate cellular chlorophyll content, specific growth rates, and carbonate chemistry. Cells were counted with a Neubauer hemacytometer chamber on a Zeiss inverted light microscope. Chlorophyll was measured on a Turner fluorometer following overnight extraction in 100% methanol at-20°C. Relative fluorescence units were recorded before and after the addition of two drops 0.1 N HCl and total chlorophyll was calculated by the method described in Holm-Hansen *et al*. 1965^38^. *B. nutricula* chlorophyll was measured using a Multiskan SkyHigh Microplate UV-Vis spectrophometer (Thermo Scientific) following overnight extraction in 100% methanol at-20°C and calculated by the method described by Ritchie 2006^39^. These results are summarized in Extended Data Fig. 2a-b.

Culture pH was measured using an UltraBASIC pH meter (Denver Instruments) equipped with an Orion glass pH electrode (ThermoFisher Scientific) calibrated with NBS standard buffer. Total CO_2_ was measured using a Corning 965 Carbon Dioxide Analyzer with technical duplicates taken between 10 mM NaCO_3_ standard measurements. Carbonate chemistry in the cultures was calculated using CO_2_Sys_v2.1^40^ by inputting salinity, temperature, pH and TCO_2_.

### Transcriptomics

Transcriptomic datasets and analyses were performed as described in Yee et al. 2019^21^. In brief, the VHA mRNA expression levels Fragments Per Kilobase Million (FPKM) was acquired from transcriptomics analysis of RNAseq data collected from time-course experiments through silicon starvation and following silicon replenishment from biological duplicates of synchronized *T. pseudonana* cultures. The data discussed in this publication have been deposited in the NCBI’s Gene Expression Omnibus^41^ and are accessible through GEO Series accession number GSE75460 (https://www.ncbi.nlm.nih.gov/geo/query/acc.cgi?acc=GSE75460) and GSE203136 (https://www.ncbi.nlm.nih.gov/geo/query/acc.cgi?acc=GSE203136).

### Microscopy

*T. pseudonana* expressing VHA_B_-eGFP were incubated with the acidotropic pH stain 2-(4-pyridyl)-5-((4-(2-dimethylaminoethyl-aminocarbamoyl)methoxy)phenyl)oxazole (PDMPO) (LysoSensor YellowBlue DND-160; Life Technologies) at a final concentration of 0.125 μM without washing in F/2. Cells were then transferred to a 35 mm poly-d-lysine coated glass bottom dish and mounted on a Warner Instruments QE-1HC Quick Exchange Heated/Cooled stage chamber controlled by CL-200 Dual Channel Temperature Controller maintained at 18 °C. Cells were imaged with a Zeiss LSM800 inverted confocal microscope equipped with a Zeiss Plan Apochromat 63× (1.4) Oil DIC M27 objective, and Zeiss Airyscan super-resolution detector. Z-stacks of three channels were acquired to monitor eGFP (Ex 488 nm with 0.3% laser power, Em 509 nm, and detection 490-535 nm), PDMPO (Ex 335 nm at 0.5% laser power, Em 530 nm, detection 550-650 nm), and chlorophyll (Ex 488 nm with 0.2% laser power, Em 667 nm, detection 450-700 nm) fluorescence. 3D images were generated and exported using the Zen Blue software package (Carl Zeiss Microscopy GmbH).

### Oxygen measurements

Individual cultures of *T. pseudonana*, *P. tricornutum*, *E. huxleyi*, *C. protothecoides*, and *P. purpurneum* grown in F/2 were maintained in exponential growth in 1 l bioreactor tubes bubbled with air and under continuous illumination by cool-white fluorescent lamps at ~100 μmol photons m^-2^ s^-1^ at 18°C. Replicate cultures of *B. nutricula* grown in K2 were maintained in exponential growth in T25 tissue culture flasks under a 12h light-dark cycle illuminated by cool-white fluorescent lamps at ~70 μmol photons m^-2^ s^-1^ at 2Ü°C. Cultures were sampled over multiple days to obtain replicate measurements of oxygen production rates.

Samples were pre-incubated with vehicle DMSO (control) or 10 nM concanamycin A [a highly specific of VHA^42^] for 10 mins before measurement. One ml of sample was used to measure oxygen production on a Hansatech Oxy-lab Clark-type electrode operated with O2 view software (Hansatech Instruments) that controlled illumination with a red LED light at 1000 μmol photons m^-2^ sec^-1^ and stirring within the water-jacketed chamber maintained at 18°C. For *B. nutricula*, 2 ml of sample was used to measure oxygen production with an optical oxygen sensor operated with Presens Measurement Studio 2.0 (PreSens Precision Sensing GmbH) and illuminated with a bright white LED light at 500 μmol photons m^-2^ sec^-1^ and stirring at room temperature (~20°C). Net maximum O_2_ production was measured by calculating the slope of O_2_ concentration over ~10 mins of illumination. Respiration was calculated similarly over 1 min immediately after illumination.

Gross maximum production was calculated by the equation:

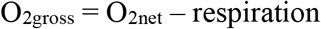

O_2_ production rates were analyzed in Graphpad Prism 6 (GraphPad Software, La Jolla California USA) by paired two-tailed t-test between control and concanamycin A treatments for each species. Oxygen production rates and P-values showing statistical significance between paired treatments are summarized in Extended Data Fig. 2c and Extended Data Table 1.

### Nanoscale secondary ion mass spectrometry (NanoSIMS)

*T. pseudonana* cells grown in F/2 were harvested from 1l tube bioreactors at ~1.6×10^6^ cells/ml. Cells were washed and resuspended in artificial seawater (ASW) medium^43^ at ~1×10^6^ cells/mL for stable isotope labelling. Incubations were carried out in 20 ml of culture with 2 mM NaH^13^CO_3_ and vehicle DMSO or 10 nM concanamycin A (dissolved in DMSO) in 50 ml Erlenmeyer flasks in duplicate at 18°C under ~500 μmol photons m^-2^ s^-1^ for 8h followed by addition of paraformaldehyde (3.7% final conc.). Controls included a culture without NaH^13^CO_3_ (but with unlabeled NaHCO_3_) and a second culture where cells were fixed for 30 min and 2 mM NaH^13^CO_3_ subsequently added. All samples were pelleted and resuspended in silica-free ASW+10% hydrofluoric acid on a rotating incubator overnight to strip the cell wall in order to eliminate charging of Si during SIMS analysis. Triplicate washes in silica-free ASW were used to remove the acid before storage at 4°C.

One ml of sample was filtered onto 0.2 μm pore size white polycarbonate filters (Millipore, Billerica, MA). After drying, filters were cut into ~1/8 sections and placed onto an analysis bullet using adhesive and conductive tabs (#16084–6, Ted Pella, Redding, California) and sputter coated with ~5 nm gold particles. A Cameca NanoSIMS 50 at Lawrence Livermore National Laboratory generated isotope images by rastering the primary ^133^Cs^+^ ion beam (2 pA, ~150 nm diameter, 16 keV) across 35 × 35 μm analysis areas with 256 × 256 pixels and a dwell time of 1 ms/pixel, for 25 scans (cycles). Prior to ion collection, analysis areas were sputtered with 90 pA of Cs+ current to a depth of ~60 nm to achieve sputtering equilibrium and optimal ionization of intracellular isotopic material. Mass resolving power of the secondary ion mass spectrometer was tuned for ~7000 (corrected units) to optimally collect quantitative secondary ion images for ^12^C^12^C– and ^13^C^12^C-on individual electron multipliers in pulse counting mode, as previously described^44^.

Image data generated from the NanoSIMS were processed and analyzed using L’Image (http://limagesoftware.net). Following dead-time and image shift correction, ^13^C^12^C-/^12^C^12^C-ratio images were produced, revealing the level and location of ^13^C fixation into biomass from H^13^CO_3_^−^ in *T. pseudonana* cells treated with DMSO or concanamycin A. Regions of interest (ROIs) were drawn around cells based on their ^12^C^14^N- and δ-^13^C enrichment, which together indicated the centric biomass shape of each cell (Extended Data Fig. 5). From each ROI, the ^13^C^12^C-/^12^C^12^C– ratios were extracted by cycle and averaged and then divided by two to calculate the ^13^C/^12^C ratio^45^. The ^13^C/^12^C ratios were then used to calculate net algal biomass produced from C fixation over the 8 h incubation period for each cell, called C_net_^44^.

The NanoSIMS data were analyzed with Graphpad Prism 6 software and in RStudio. Values were checked for normality using Kolmogorov-Smirnov tests. Statistical comparisons of C-fixation data were performed with unpaired t-test with Welch’s correction to compensate for different standard deviations between the controls and treatment.

### ^14^C photosynthesis versus irradiance curves

Cultures of *T. pseudonana* grown in F/2 in 1 l bioreactor tubes bubbled with air and under continuous illumination by cool-white fluorescent lamps at ~100 μmol photons m^-2^ s^-1^ at 18°C. The control (with vehicle DMSO) and 10 nM concanamycin A treated P-E incubations were performed over three days of batch culturing at standard DIC and low DIC relative to seawater levels (Extended Data Fig. 4b). Standard DIC conditions are at levels from unmodified culture medium while low DIC was reached by titrating cultures with 1M HCl and bubbling off excess CO_2_ with air just prior to the incubations.

Photosynthesis-Irradiance (P-E) incubations were carried out by incubating cells in a modified ^14^C bicarbonate incorporation technique described by Lewis & Smith 1983^46^. Experiments were carried out in four custom P-E incubators maintained at 18°C and at 18 discreet irradiances ranging from 0 to 1678 μmol photons m^-2^ sec^-1^ illuminated by a 150W tungsten-halogen lamp. Discreet measurements of total photosynthetically available radiation (PAR, 400-700 nm) were taken with a Spherical Micro Quantum Sensor US-SQS (Heinz Walz, GmBH). For each P-E incubation, 100 μl of a 1 mCi NaH^14^CO_3_(Perkin-Elmer) solution was added to 25 ml of diatom culture samples to reach a final concentration of 4 μCi ml^-1^. Spiked culture was aliquoted in 1 ml volumes into 21 pre-chilled 7 ml borosilicate glass scintillation vials. Eighteen of these vials were illuminated for 1 h in the P-E incubator while three remaining vials were immediately acidified with 150 μl of 12N HCl to drive off inorganic carbon and determine background activity for time-zero (T0) samples. Three total radioactivity (TA) samples were generated by combining 50 μl of spiked culture with 200 μl of phenylethylamine (Sigma-Aldrich) topped off with 5 ml of Ecolite(+) scintillation cocktail (MP Biomedicals) and capped immediately. Incubations were terminated after 1 h by switching off the lamps and adding 150 μl of 12N HCl to each vial. All acidified samples were allowed to exhaust un-fixed DIC overnight in a fume hood and topped off with 5 mL of Ecolite(+) and capped the next day for counting radioactivity on a Beckman Coulter LS6500 liquid scintillation counter.

Bulk carbon fixation rates (*P_bulk_*; mg C l^-1^ hr^-1^) were calculated from disintegrations per minute (DPM) by the equation:

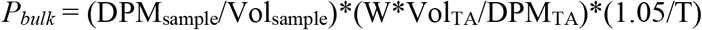

Where DPM_sample_ is the T0 corrected activity per sample, Vol_sample_ is volume of culture per sample, W is the DIC concentration of the culture, Vol_TA_ is volume of culture in TA samples, DPM_TA_ is the activity per TA sample, 1.05 is the discrimination factor between ^14^C and ^12^C uptake, and T is the time duration of the incubations.

### P-E curve fitting and calculation of VHA contribution

*P_bulk_* was normalized by cell density to give cellular carbon fixation rates (*P*; fmol C cell^-1^ min^-1^) and 18-point curves were fit with the equations by Platt et al. 1980^47^ using the Phytotools R-package (https://CRAN.R-project.org/package=phytotools). Fitted curves were plotted from 0-1600 μmol photons m^-2^ sec^-1^ and averaged amongst the respective conditions (Extended Data Fig. 4a).

Paired two-tailed t-test were carried out on averages of the curve fitting parameters *Pmax*(maximum cellular carbon fixation rate), *α* (initial slope), and *E_k_* (half-saturation constant of photosynthesis) between control and concanamycin A treatments using Graphpad Prism 6. P-values showing statistical significance for each parameter are summarized in Extended Data Table 2.

The contribution of VHA activity to *P* was calculated between control and concanamycin A treated average curve fits by the following equation:

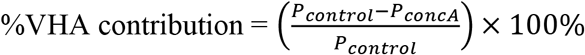

### Data Availability

The data discussed in this publication have been deposited in the NCBI’s Gene Expression Omnibus and are accessible through GEO Series accession number GSE75460 (https://www.ncbi.nlm.nih.gov/geo/query/acc.cgi?acc=GSE75460) and GSE203136 (https://www.ncbi.nlm.nih.gov/geo/query/acc.cgi?acc=GSE203136).

## Acknowledgments

We are grateful for the help of Dr. Roshan Shreshta, Dr. Brian Palenik, Dr. Orna Cook, Dr. Christopher Hewes, Dr. Garfield Kwan, Dr. Eric Armstrong and Mr. Angus Thies (details in Suppl. Text). Work at LLNL was performed under the auspices of US Department of Energy Office of Science contract DE-AC52-07NA27344. The confocal microscope at the Tresguerres laboratory was generously donated by the Arthur M. and Kate E. Tode Research Endowment in Marine Biological Sciences (UCSD).

## Author contributions

Conceptualization: DPY, MH, MT

Methodology: DPY, TJS, RMA, MV, XM, PKW, BGM, MH, MT

Investigation: DPY, TJS, BS, MH, MT

Visualization: DPY, TJS, MT

Funding acquisition: DPY, TJS, XM, PKW, MT

Project administration: DPY, MT

Resources: MH, MV, XM, BGM, MT

Supervision: MT

Writing – original draft: DPY, MT

Writing – review & editing: DPY, TJS, RMA, MV, PKW, XM, MT

Authors declare that they have no competing interests.

Supplementary Information is available for this paper.

Correspondence and requests for materials should be addressed to Daniel P. Yee or Martin Tresguerres.

Reprints and permissions information is available at www.nature.com/reprints.

## Extended Data

**Extended Data Figure 1.**
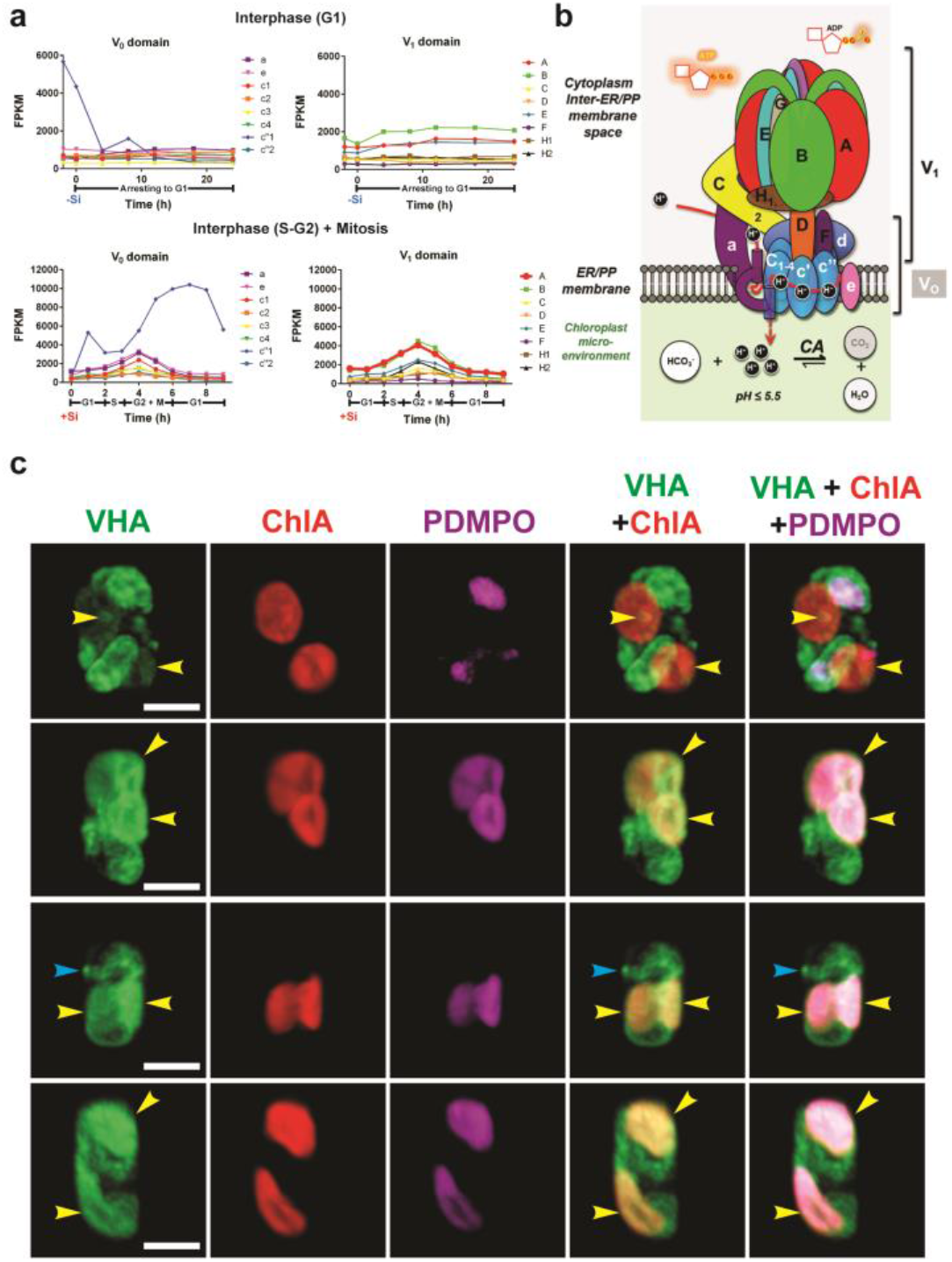
Expression and subcellular localization of VHA in *T. pseudonana*. (**a**) Transcriptomic profile of VHA subunits during G1 (cell-cycle arrest following silicon starvation; top), and G2-S-G2 mitosis (synchronous division following silicon readdition; bottom). (**b**) Diagram of the VHA-holoenzyme complex in the endoplasmic reticulum/periplastid (ER/PP) membranes surrounding the chloroplast. CA = carbonic anhydrase. (**c**) 3D confocal images of eGFP-tagged VHAB around chloroplasts (yellow arrows) co-localized with the acidotrophic dye PDMPO (magenta), and silica deposition vesicles (blue arrows) at different cellcycle stages [scale bars: 5 μm].

**Extended Data Figure 2.**
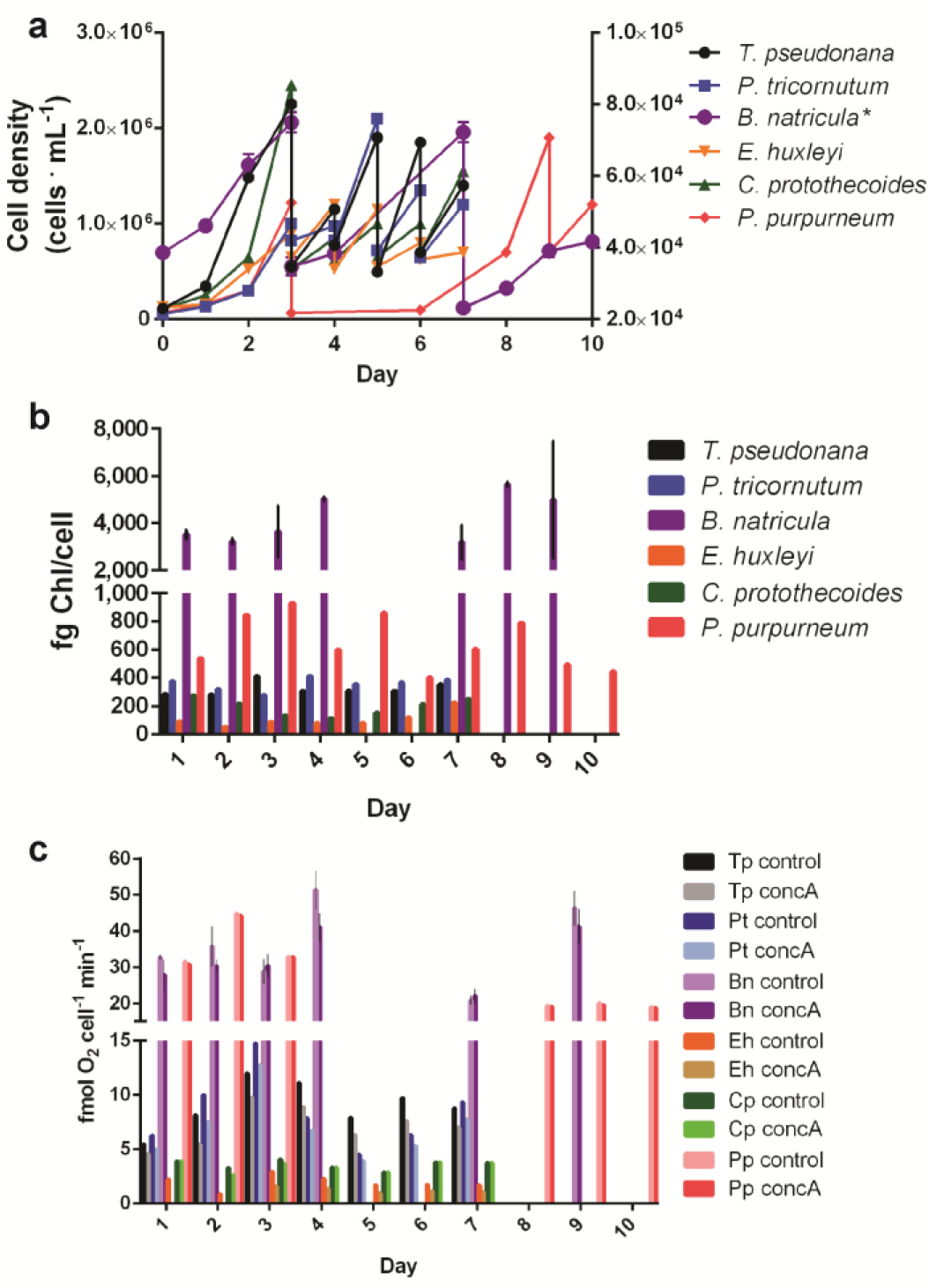
Daily culture and O_2_ production measurements for six marine algae. (**A**) Growth curves, (**B**) chlorophyll content per cell, and (**C**) gross maximum O_2_ production per cell in control (vehicle DMSO) and VHA-inhibited (concA=10 nM concanamycin A). Tp=*T. pseudonana* (centric diatom); Pt=*P. tricornutum* (pennate diatom); Bn=*B. nutricula* (dinoflagellate; n=2 before & 3 after day 7); Eh=*E. huxleyi* (coccolitophorid); Cp=*C. protothecoides* (green algae); Pp=*P. purpurneum* (red algae); *denotes data assigned to the right axis.

**Extended Data Figure 3.**
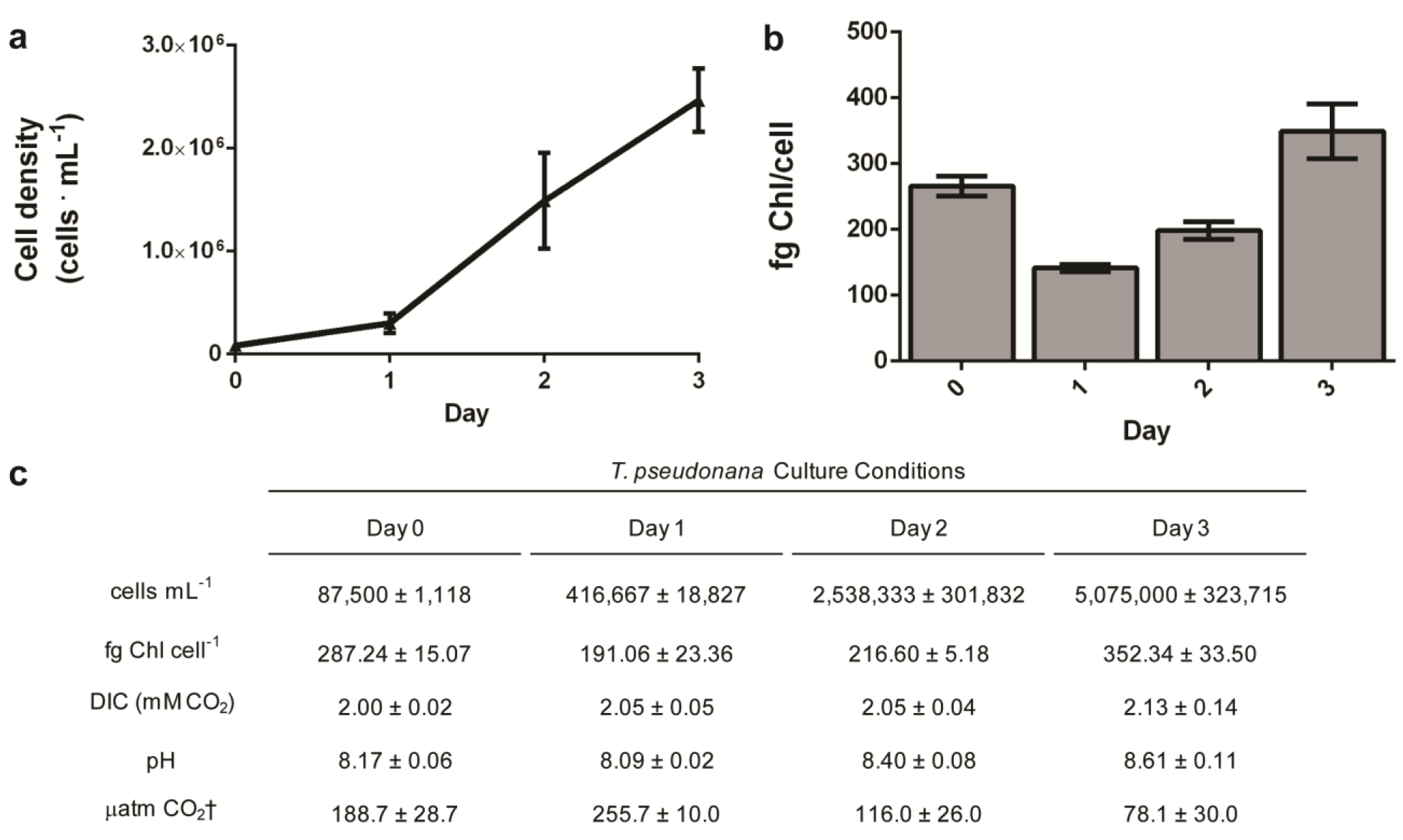
Culture conditions for *T. pseudonana* used in ^14^C P-E experiments. (**a**) cell growth, (**b**) chlorophyll content per cell, and **(c)**summary of daily measurements of cells mL^-1^, fg Chl ml^-1^, DIC, pH, and pCO_2_ (n=6, error bars= SEM, ^†^denotes values calculated using CO2Sys).

**Extended Data Figure 4.**
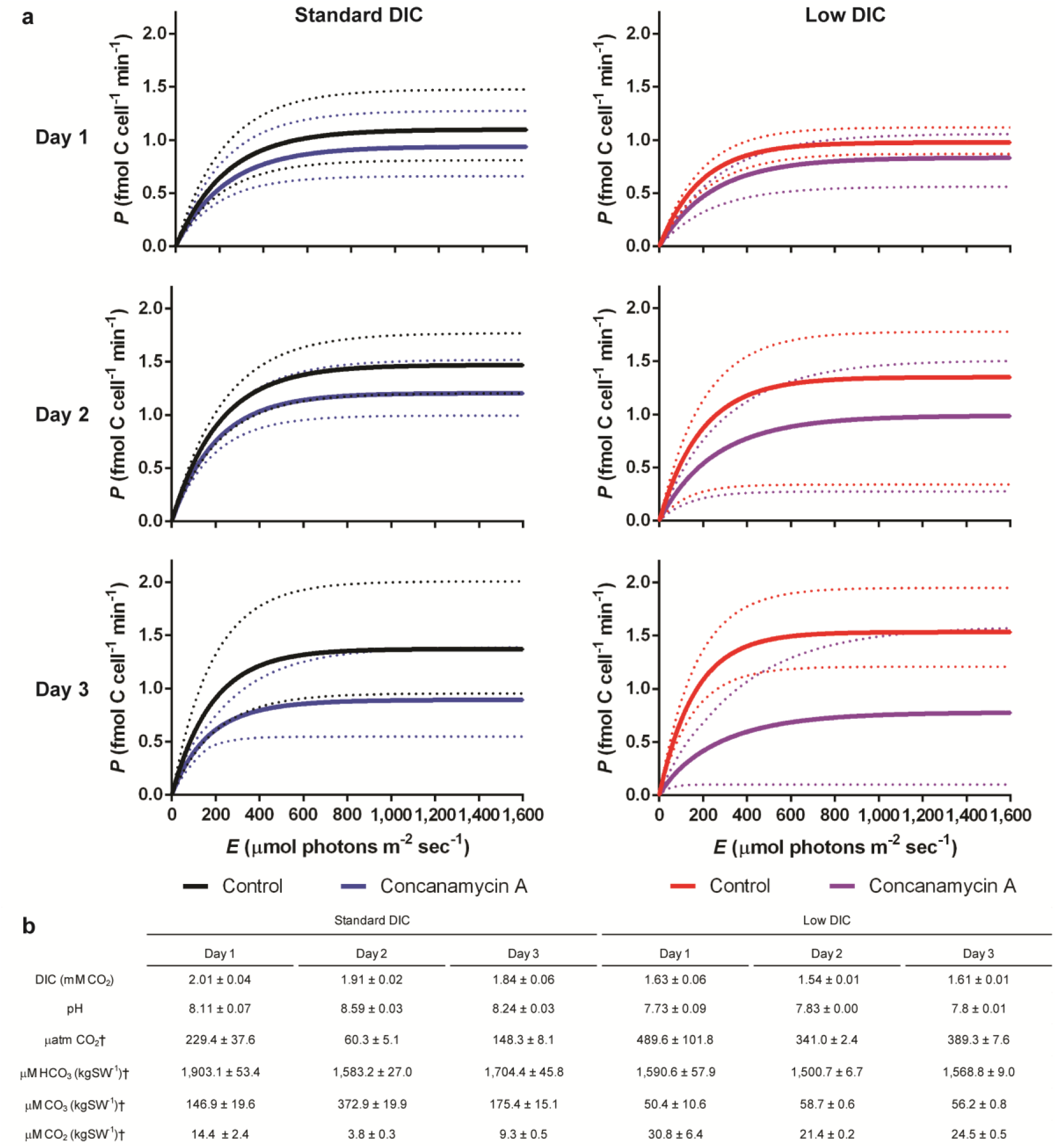
Photosynthetic carbon fixation by *T. pseudonana*. (**a**) ^14^C P-E curves of control (vehicle DMSO) and VHA-inhibited (10 nM concanamycin A) fitted from 0-1,600 μmols photons m^-2^ sec^-1^ measured over three days of culturing under standard and low DIC conditions (n=6, dotted lines= range). (**b**) Table summarizing the carbon chemistry conditions from respective ^14^C P-E incubations (n=3, ^†^denotes values calculated using CO2Sys).

**Extended Data Figure 5.**
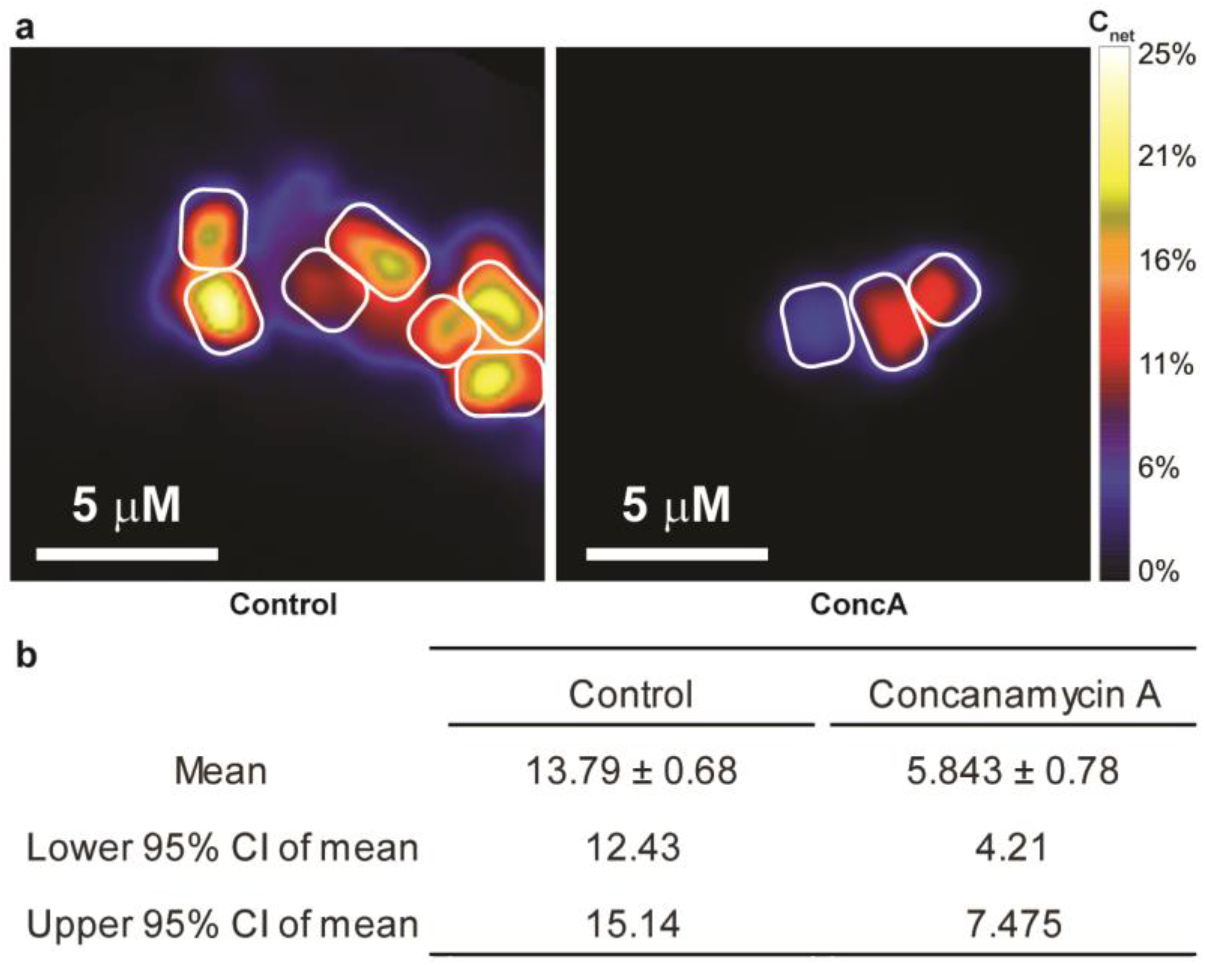
Single-cell NanoSIMS quantification ^13^C-fixation (a)representative. NanoSIMS images of *T. pseudonana*, Enrichment of ^13^C in control (left) and concanamycin A (right) treated cells used for single-cell quantification of net biomass formation from photosynthesis, **(b)**statistics for quantification of net biomass formation from photosynthesis (Fig. 3a)

**Extended Data Table 1.** Values and statistics from oxygen production measurements.

|  | fmol O<sub>2</sub> cell<sup>-1</sup> min<sup>-1</sup> |  |  | % O<sub>2</sub> Production |
| --- | --- | --- | --- | --- |
|  | Control | ConcA | P-value | Mean |
| *T. pseudonana* | 8.79 ± 0.53 | 6.96 ± 0.36 | 0.0090** | 20.74 ± 0.74 |
| *P. tricornutum* | 7.82 ± 0.88 | 6.58 ± 0.70 | 0.0203* | 15.80 ± 0.62 |
| *B. natricula* | 44.75 ± 3.88 | 36.14 ± 3.59 | 0.0152* | 19.14 ± 4.37 |
| *E. huxleyi* | 1.92 ± 0.19 | 1.17 ± 0.12 | 0.0125* | 39.16 ± 2.20 |
| *Chlorella* sp. | 3.47 ± 0.30 | 3.47 ± 0.30 | nd | 0.00 |
| *P. purpureum* | 19.39 ± 0.32 | 19.09 ± 0.21 | 0.122 | 1.55 ± 0.57 |

Mean ± SEM gross maximum O<sub>2</sub> production, and % O<sub>2</sub> reduction of six microalgae species along with their P-values between paired control and concanamycin A treatments (nd= no difference).

**Extended Data Table 2.** Curve fitting parameters and statistics from ^14^C P-E incubations.

| Standard DIC |  |  |  |  |  |  |  |  |  |
| --- | --- | --- | --- | --- | --- | --- | --- | --- | --- |
|  | Control |  |  | Concanamycin A |  |  | P-value |  |  |
|  | Day 1 | Day 2 | Day 3 | Day 1 | Day 2 | Day 3 | Day 1 | Day 2 | Day 3 |
| $P_{max}$ | 1.1 ± 0.09 | 1.47 ± 0.11 | 1.37 ± 0.19 | 0.94 ± 0.08 | 1.21 ± 0.08 | 0.89 ± 0.12 | < 0.0001**** | 0.017* | 0.0037** |
| $a$ | 0.0048 ± 0.0004 | 0.0069 ± 0.0003 | 0.0075 ± 0.0009 | 0.0041 ± 0.0004 | 0.0059 ± 0.0003 | 0.0054 ± 0.0005 | 0.0019** | 0.0013** | 0.0485* |
| $E_k$ | 227.03 ± 4.49 | 213.13 ± 8.73 | 182.61 ± 7.01 | 230.01 ± 10.68 | 202.93 ± 8.58 | 167.37 ± 22.43 | 0.7074 | 0.2917 | 0.5376 |
| $n$ | 6 | 6 | 6 | 6 | 6 | 6 | | | |
| Low DIC |  |  |  |  |  |  |  |  |  |
|  | Control |  |  | Concanamycin A |  |  | P-value |  |  |
|  | Day 1 | Day 2 | Day 3 | Day 1 | Day 2 | Day 3 | Day 1 | Day 2 | Day 3 |
| $P_{max}$ | 0.98 ± 0.04 | 1.35 ± 0.28 | 1.53 ± 0.16 | 0.83 ± 0.08 | 0.99 ± 0.26 | 0.78 ± 0.27 | 0.0351* | 0.0347* | 0.0075** |
| $a$ | 0.0052 ± 0.0003 | 0.0071 ± 0.0012 | 0.0094 ± 0.0008 | 0.0035 ± 0.0003 | 0.0041 ± 0.0007 | 0.0036 ± 0.0005 | 0.005** | 0.0122* | 0.0499* |
| $E_k$ | 191.29 ± 8.06 | 179.73 ± 15.63 | 161.93 ± 3.87 | 241.39 ± 17.80 | 217.05 ± 35.04 | 185.47 ± 56.35 | 0.0039** | 0.0133* | 0.098 |
| $n$ | 5 | 4 | 3 | 5 | 5 | 5 | | | |

Mean ± SEM of $P_{max}$ (maximum cellular carbon fixation rate; fmol C cell<sup>-1</sup> min<sup>-1</sup>), $a$ (initial slope), and $E_k$ (half-saturation constant of photosynthesis; μmol photons m<sup>2</sup> sec<sup>-1</sup>), along with corresponding $n$ values. P-values of curve fitting parameters between paired control and concanamycin A treatments.

**Extended Data Table 3.** Estimates of VHA contribution to primary production *(PP)*

|  | Gigatons C year<sup>-1</sup> | Contribution from diatoms | Contribution from diatom VHA† | Gigatons C year<sup>-1</sup> from diatoms |
| --- | --- | --- | --- | --- |
| Marine PP‡ | 53.5 | 48.6% | 6.6 - 25.3% |  |
| Global PP‡ | 115.6 | 22.5% | 3.0 - 11.7% | 3.5-13.5 |
| Diatom PP§ | 26.0 | 100% | 13.5 - 52% |  |

‡From Field *et al.* 1998, §From Nelson *et al.* 1995, †Estimated based on VHA contribution to diatom photosynthesis ranging from 13.5% to 52% (Fig. 3, this study).

## Supplementary Information

### Supplementary Notes

#### Extended acknowledgments

Dr. Roshan Shreshta provided plasmids and assistance with cloning and transformations. Dr. Orna Cook, Dr. Johan Decelle and Dr. Charlotte Le kieffre provided microalgae cultures. Dr. Christopher Hewes assisted with fluorometric chlorophyll measurements. Dr. Brian Palenik and Dr. Giovanni Finazzi provided oxygen measurement equipment. Dr. Garfield Kwan, Angus Thies and Dr. Eric Armstrong provided images for Fig. 4. The Arthur M. and Kate E. Tode Research Endowment in Marine Biological Sciences (UCSD) provided essential equipment used in this project.

#### Funding and Grant Numbers

National Institutes of Health grant T32GM067550 (DPY)

Ralph Lewin Graduate Fellowship (DPY)

LLNL Laboratory Directed Research and Development (LDRD) project # 19-LW-044

(TJS, XM)

SIO discretionary funds (MT)

